# Planting 10 Million Trees with Lukenya University in Kenya: Methodology and Preliminary Observations and Forecasts

**DOI:** 10.1101/2025.01.09.632148

**Authors:** Protus Kyalo, Vyacheslav Kungurtsev, Esther Muli, Stephan Pietsch, Kennedy Mugo, Margrate Kaigongi, Martin Muasa, Judith Wafula, Sammy Muvela, Charles Mutua

## Abstract

Lukenya University has recently begun implementing a Ten Million Tree Growing Initiative, as part a larger national multi-institutional country-wide tree growing program. In this paper we describe the methods used as far as tree selection and the planting itself. Note that most of the trees are not planted directly by the institution by given to farmers to plant on their land, with instruction as to how to choose a location and how to technically perform the planting and maintenance to ensure greater success for the tree and the environment surrounding it. This knowledge arises out of years of experience as far as both the biology of planting and the social interaction with smallholder farmer communities. We kept track of the survival and growth of a subsample of a number of different tree species. After about two years of operation, we performed a statistical analysis on preliminary data. We observe statistically significant advantages to survival and relative growth volume to indigenous trees and the aforementioned comprehensive instruction before plotting, among other observations we relate from the hypothesis tests performed. Finally, we present estimates of the carbon sequestration and economic value of the trees to the community. Summarizing, the paper presents a comprehensive transparent display of a tree growing initiative, a common endeavor motivated to maximize social welfare and climate change resilience, and in so doing can develop the knowledge of best practices for these purposes.

## 1. Introduction

Lukenya University (LU), nestled in the semi-arid landscape of Mtito Andei in Makueni County, Kenya, is an institution committed to pioneering sustainable development initiatives. Founded in 2015 and officially chartered in 2022, LU operates under a ”Green Philosophy,” integrating environmental sustainability into its core mission of education, research, and community service. This philosophy has guided the university’s efforts to address pressing environmental challenges, particularly in arid and semi-arid regions such as Kibwezi East Sub-County. Among its flagship programs is the ambitious Ten Million Trees Initiative, anchored in the Kenyan National Landscape and Ecosystem Strategy (2023-2032) that seeks to grow 15 billion trees in order to restore 10.6 million hectares of degraded land ^1^. This decade-long reforestation project aims at combating climate change, restoring ecosystems, and empowering local communities. For a recent publication see [27]. Note the formal program^2^, and a description of a marathon event to advertise the implementation of the initiative ^3^,.

The Ten Million Trees Initiative, beyond aesthetic and shading canopy benefits to the community, is expected to contribute to climate resilience, biodiversity conservation, and sustainable livelihoods. It will increase forest cover, which would provide much needed shade for farmers and nutrition for native fauna, sequester carbon and mitigate the effects of global warming, and improve the hydrological cycles that are vital for water-scarce regions. Moreover, the species chosen all contain valuable parts of the tree, including medicine, nutraceuticals, fodder for livestock, firewood, and other products, which is expected to both increase as well as improve the risk profile of smallholder income through diversification. This initiative also aligns with Kenya’s national goals for afforestation and global commitments under the United Nations Sustainable Development Goals (SDGs), particularly those related to climate action, life on land, and poverty reduction.

LU’s initiative is grounded in collaboration and community engagement. Working closely with organizations such as the Kenya Forest Service (KFS), the Kenya Forestry Research Institute (KEFRI), and international partners like UNESCO, the university has created a robust network to ensure the initiative’s success. Local farmers, youth groups, and community members are integral to this process, receiving training in sustainable forestry practices and contributing to the planting and care of the trees.

To support the initiative’s planning and implementation, LU employs advanced scientific tools and models to monitor and evaluate tree growth and survival. Among these, the Biogeochemical Management Model (BGC-MAN) [34] has emerged as a critical tool for understanding tree growth dynamics under varying hydrological conditions[35]. This model helps simulate tree responses to environmental factors such as soil type, precipitation patterns, and temperature fluctuations, providing valuable insights for optimizing tree species selection and site management. Integrating scientific innovation with traditional knowledge, the initiative aims to maximize ecological and social impact.

The Ten Million Trees Initiative also seeks to inspire and mobilize stakeholders beyond the university’s immediate community. It serves as a call to action for policymakers, environmental organizations, and the private sector to invest in scalable and sustainable solutions to environmental degradation. Through evidence-based practices and transparent reporting, LU aims to demonstrate the transformative potential of large-scale reforestation projects, attracting more partners and resources to realize the initiative’s vision. Ultimately, the Ten Million Trees Initiative reflects LU’s commitment to addressing the challenges of climate change and environmental degradation through education, research, and community action. Leveraging cutting-edge models like BGC-MAN and fostering inclusive collaborations, LU hopes not only to restore degraded lands but also to build a greener, more resilient future for generations to come. This initiative exemplifies the university’s ethos of labouring for the benefit of the future, encapsulated in its motto: Postera Crescam Laude. So far 371,162 trees have been distributed as of November, 2024.

## 2. Related Works: Reforestation Initiatives

Large-scale tree planting initiatives have emerged as crucial strategies in combating climate change and environmental degradation worldwide. The Great Green Wall initiative spanning across Africa’s Sahel region represents one of the most ambitious reforestation projects, aiming to restore 100 million hectares of degraded land [37]. This initiative has demonstrated both the potential and challenges of large-scale restoration efforts, particularly in arid and semi-arid regions similar to Makueni County [16].

In East Africa, Ethiopia’s Green Legacy Initiative launched in 2019 has set remarkable precedents in mass tree planting campaigns [40]. The initiative reported planting over 350 million trees in a single day, showcasing the potential of mobilizing communities for environmental conservation. However, studies have highlighted that success in such initiatives depends heavily on post-planting care and community engagement rather than initial planting numbers [4].

Kenya’s own history with large-scale reforestation provides valuable lessons for current initiatives. The Green Belt Movement, founded by Nobel laureate Wangari Maathai, has planted over 51 million trees since its inception in 1977 [25]. Their community-based approach to tree planting has become a model for sustainable reforestation, emphasizing the importance of indigenous species and local community participation in environmental conservation efforts [20].

Research on reforestation in semi-arid regions has highlighted several critical factors for success. Studies in similar ecological zones have shown that species selection, timing of planting, and post-planting care significantly influence survival rates [13]. The use of indigenous species, in particular, has been found to result in higher survival rates and better ecosystem integration compared to exotic species in arid environments [6].

Recent technological innovations have enhanced the potential for successful reforestation projects. The integration of geographic information systems (GIS) and remote sensing technologies has improved site selection and monitoring capabilities [21]. Additionally, advances in modelling tools, such as the BGC-MAN model, have enabled better prediction of tree growth patterns and survival rates in challenging environments [7].

Corporate and institutional involvement in reforestation has also evolved significantly. Universities worldwide have increasingly taken leadership roles in environmental conservation, with many institutions developing comprehensive sustainability plans that include tree planting components [19]. These academic initiatives often combine research, education, and community outreach, similar to Lukenya University’s approach.

The success of reforestation initiatives in Kenya has been significantly influenced by policy frameworks and institutional support [2]. The Kenya Forest Service’s Farm Forestry Program has demonstrated the importance of integrating tree planting with agricultural practices, particularly in arid and semi-arid regions [30]. This approach has shown promise in addressing both environmental conservation and food security concerns.

Global experiences with large-scale reforestation have provided valuable insights into best practices. China’s Grain for Green Program, one of the world’s largest ecological restoration initiatives, has shown that successful reforestation requires long-term commitment, substantial resource allocation, and strong policy support [9]. Similarly, Costa Rica’s national reforestation program has demonstrated the effectiveness of incentive-based approaches in promoting forest conservation and restoration [8].

Monitoring and evaluation frameworks have become increasingly sophisticated in recent years. Satellite imagery and drone technology are now commonly used to track planting progress and forest cover changes [24]. These technological advancements, combined with traditional ground-based monitoring methods, provide more accurate assessments of reforestation success rates and impact [18].

### 2.1. Contributions

In this paper, we make several contributions to the scientific literature:

- We provide a description of the planting techniques utilized as part of the initiative, and the means by which they are taught to farmers. We describe broad insights on the most succcessful practices as well as innovations developed by Lukenya.
- We give a description of the species chosen, including what they provide to the community.
- We present a statistical analysis on preliminary data on tree survival rate, presenting the results for statistical tests to ascertain patterns.
- We extrapolate an estimate for the total carbon sequestered, and the total economic valuation of all the products produced by the initiative.

## 3. Native Flora of East Africa

### 3.1. Landscape and Eco-zones of Kenya

Kenya’s landscape is characterized by remarkable diversity, ranging from coastal regions to highland plateaus and arid lowlands. The country’s topography is dominated by the Great Rift Valley, which bisects the western highlands, creating distinct ecological zones that support varied vegetation patterns [23]. These highlands, including Mount Kenya and the Aberdare Range, receive significant rainfall and historically supported extensive forest coverage.

The country is divided into seven major ecological zones, from humid highland forests to arid and semi-arid lowlands. This ecological gradient is strongly influenced by altitude and rainfall patterns, with annual precipitation varying from over 2,000mm in the highlands to less than 250mm in the northern arid regions [28] [3]. These variations create distinct vegetation zones, each supporting unique plant communities adapted to local conditions.

### 3.2. Common Plant Families in Kenya

Kenya’s flora is represented by approximately 7,000 plant species, with the most prevalent families including *Fabaceae* (legumes), *Asteraceae* (daisies), and *Poaceae* (grasses). The *Euphorbiaceae* family is particularly well-adapted to arid conditions, while the *Rubiaceae* family dominates in forest understories [33]. These plant families have evolved diverse adaptations to survive in Kenya’s varied ecological conditions, from drought-resistant characteristics to specialized pollination mechanisms.

Indigenous tree species such as Acacia (now *Vachellia* and *Senegalia*), *Combretum*, and *Commiphora* are widespread across Kenya’s drylands, while highland areas feature prominent species from the *Podocarpaceae* and *Oleaceae* families [17]. Many of these native species play crucial roles in traditional medicine, provide essential ecosystem services, and support local livelihoods through various other means.

### 3.3. History of Deforestation and Reforestation

Kenya’s forest cover has experienced significant decline since the colonial period, dropping from an estimated 30% in the early 1900s to less than 6% by 2000. Major drivers of deforestation include agricultural expansion, urban development, and unsustainable charcoal production [29]. The loss of indigenous forests has had severe implications for biodiversity, water resources, and local communities dependent on forest resources.

**Figure 1:**
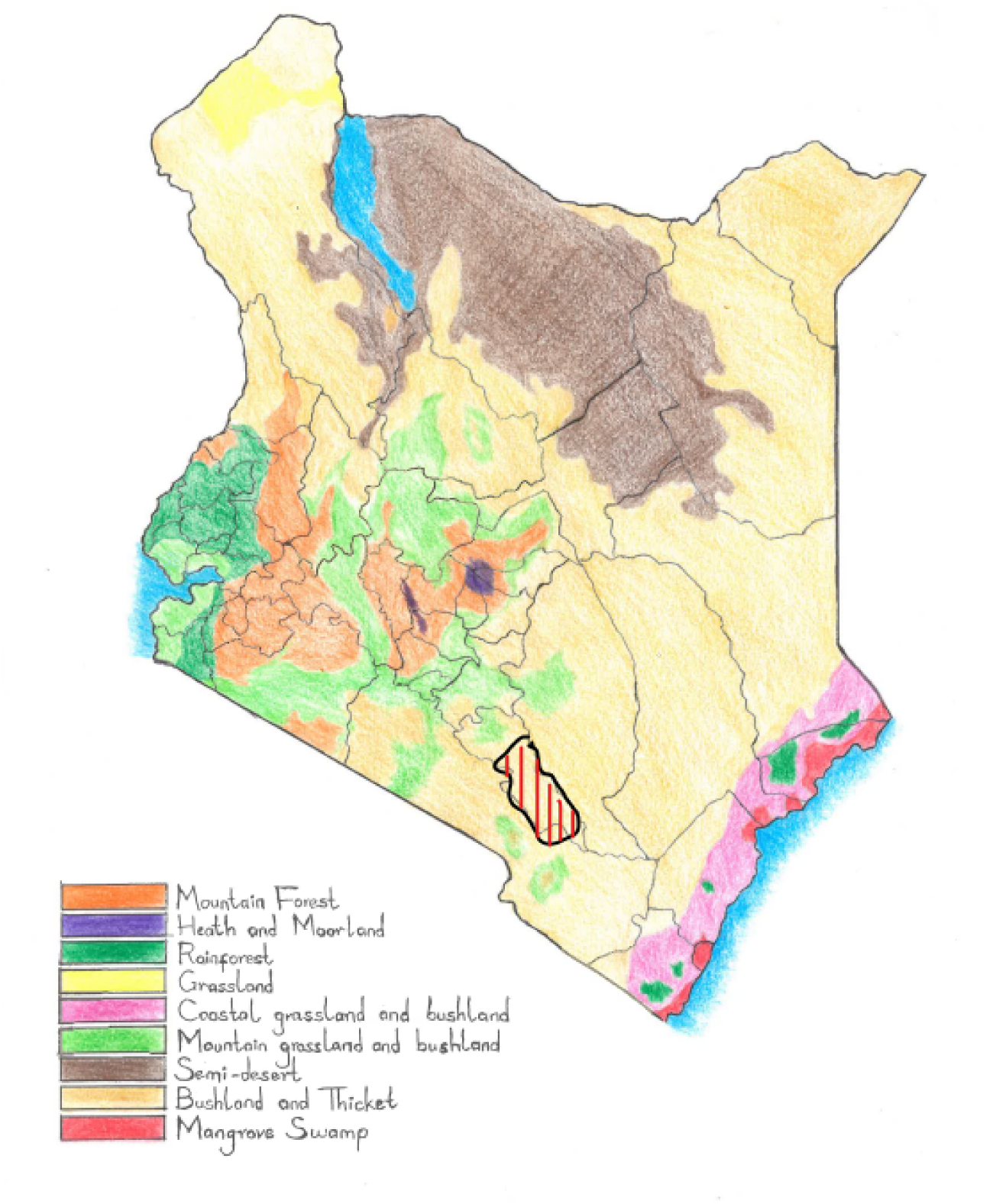
All Ecological Zones in Kenya. The dashed red lines with the black exterior in the South is the region wherein the trees discussed in this paper were planted.

Reforestation efforts in Kenya gained momentum in the 1970s with initiatives like the Green Belt Movement and government-led programs. Despite these efforts, challenges persist in balancing conservation with development needs. Recent national initiatives aim to achieve 10% forest cover by 2030, focusing on both exotic and indigenous species for restoration [22]. Success stories from community-based forest management programs demonstrate the potential for sustainable forest restoration when local communities are actively involved in conservation efforts

## 4. Tree Growing Initiative: Details

### 4.1. Chosen Species

The tree species selection for Lukenya University’s Ten Million Trees Initiative reflects a careful consideration of the semi-arid conditions in Kibwezi East, Makueni County. The selected species encompass both indigenous and naturalized varieties, chosen for their adaptability to local climate conditions and their multiple socio-economic benefits. The species can be categorized into several functional groups based on their primary uses and ecological roles.

#### 4.1.1. Fruit Trees for Food Security and Income Generation

Several fruit-bearing species have been prioritized in the initiative, combining nutritional value with economic potential. These include:

1. *Mangifera indica* (Mango/Muembe): Provides fruits and shade
2. *Citrus limon* (Lemon/Mdimu): Valued for fruits and shade
3. *Citrus sinensis* (Sweet Lemon/Chungwa): Offers fruits and shade coverage
4. *Persea americana* (Avocado): Provides nutritious fruits and canopy cover
5. *Carica papaya* (Pawpaw/Papai): Important for fruit production
6. *Psidium guajava* (Guava): Offers both nutritional and medicinal value
7. *Tamarindus indica* (Tamarind/Ukwaju): Multi-purpose tree providing fruits, medicine, and shade.
8. *Sclerocarya birrea* (Marula): Multipurpose tree utilised for fruits, seed oils, shade and medicine.
9. *Berchemia discolor* (bird plum/ brown ivory): A multipurpose tree planted for fruits, fuelwood, medicine, apiculture and shade.
10. *Thespesia garckeana* (African chewing gum/ wild hibiscus): Grown for Fruits and medicinal uses.
11. *Vangueria madagascariensis* (Spanish-tamarind): Used for fruits and shade.
12. *Vitex payos* (Chocolate berry): A multipurpose tree providing fruits, fuelwood, timber and medicine.
13. *Maerua decumbens* (Blue bush-cherry ): A multipurpose tree that is utilized for fruits, water purification, medicine, fodder and bee forage.
14. *Diospyros scabra* (Hard-leaved monkey plum): A widespread tree used for fruits, timber, medicine, fuelwood, fodder and shade.
15. *Balanites aegyptiaca* (Desert dates): A tree used for fruits, medicine and shade.
16. *Adansonia digitata* (Baobab/ Mbuyu): A multipurpose tree that provides, fruits, shade, medicine and fiber.
17. *Cordia sinensis* (Grey-leaved cordia): Valued for fruits, timber, medicine, fuelwood, shade, bee forage and fodder.

#### 4.1.2. Medicinal and Multi-purpose Trees

The initiative includes several species with significant medicinal value:

1. *Azadirachta indica* (Neem/Mwarubaini): Known for extensive medicinal properties and environmental benefits
2. *Prunus sinesis* (Red Stink Wood/Mumba Aume): Valued for medicinal properties and shade
3. *Kigelia africana* (Sausage Tree/Kiatine): Traditional medicinal uses
4. *Moringa oleifera* (Moringa): Highly nutritious with multiple medicinal applications
5. *Aloe secundiflora and Aloe barbadensis*: Important medicinal species adapted to arid conditions
6. *Melia volkensii* (Mukau): Valuable for timber and windbreak functions
7. *Gliricidia sepium* (Mexican Lilac/Mumauta): Multi-functional species providing medicine, fodder, fencing, and soil improvement
8. *Terminalia spinosa* (Mwanga/ Mutula): Effective for medicine, fuel-wood, timber, fodder and live fencing
9. *Senna siamea* (Mchora): Suitable for medicine, windbreaks and shade provision
10. *Combretum schumannii* (Sand bushwillow): Used for medicine, fuel-wood, timber posts, tool handles and carving
11. *Mystroxylon aethiopicum*(Kooboo-berry): Used for medicine, dye and fencing
12. *Newtonia hildebrandtii* (Lebombo wattle): Important species for medicine and fuelwood
13. *Tabernaemontana stapfiana* (Soccerball fruit) Popularly known for medicinal uses
14. *Cordia africana* (East African cordia): Valued for medicine, timber, bee forage, shade, ornamental, mulch and soil conservation
15. *Commiphora eminii* (Kiliva): Commonly used as medicine, firewood, timber (furniture), shade, soil conservation, resin, live fence
16. *Brachylaena huillensis*(silver oak): Applied for medicinal, wood carving, fuelwood and ornamental uses
17. *Bridelia taitensis*(Bridelia): Medicine, timber, shade, mulch and riverbank stabilization
18. *Croton dichogamous* (Orange Leaved Croton): A small tree used for medicine, food flavouring and fuelwood
19. *Haplocoelum foliolosum* (Northern galla-plum): Used in traditional medicine, construction and apiculture
20. *Anisotes Ukambensis* (Anisotes): Kenyan endemic species utilized as medicine, fodder and ornamental
21. *Vepris Simplicifolia* (Vepris): A small tree widely exploited for medicinal purpose
22. *Euclea divinorum* (Diamond-leaved euclea): Widely exploited for medicinal u
23. *Acacia tortilis* (Mwaa): Adapted to local conditions, providing multiple benefits
24. *Croton megalocarpus* (Muthulu): Important indigenous species with medicinal properties
25. *Delonix elata* (Creamy Peacock/Muange): Indigenous species with multiple benefits

This diverse selection of species reflects a balanced approach to meeting both ecological and socio-economic objectives. The species mix addresses various community needs including food security, income generation, medicinal resources, and environmental services while ensuring adaptation to the local semi-arid conditions of Makueni County.

### 4.2. Cultivation Details and Distribution Strategy

#### 4.2.1. Distribution Framework

Lukenya University has implemented a dual-approach strategy for the Ten Million Trees Initiative. The first component involves direct planting within the university’s designated blocks and sections, serving as demonstration sites and research areas. The second, more extensive component focuses on community distribution, primarily targeting farmers and residents in Kibwezi East, Makueni County, with plans for expansion to neighboring counties over the initiative’s ten-year timeframe.

#### 4.2.2. Community Engagement and Education

The initiative employs a comprehensive educational approach to prepare farmers before distributing seedlings. Farmers receive detailed instruction on various critical aspects to ensure the success of tree planting efforts. These include understanding regional climatic conditions and their implications for tree survival, selecting high-value tree species suitable for both environmental conservation and economic benefits, and integrating tree planting with livestock management, particularly significant for the many pastoralist participants. Additionally, farmers are educated on irrigation requirements and effective water management techniques tailored to semi-arid conditions, as well as conservation considerations, especially given the proximity to Tsavo East and West National Parks. This thorough preparation aims to equip farmers with the knowledge and skills necessary for the sustainable integration of tree planting into their agricultural practices.

#### 4.2.3. Technical Innovation and Support

A significant advancement in the implementation of the initiative has been the development of the Geo Hydro Deep Root Technique, later evolved into the Z-U Deep Root Technique (ZUDROT), an innovative water conservation method initially formulated by Mr. Bernard Kivyatu, a PhD student in Agriculture at Lukenya University. The technique involves creating a central deeper pit (3-4 ft.) surrounded by four relatively shallower pits (2.5 ft.) where trees are planted, enhancing deep plant root development and therefore greater resilience against drought.This technique has undergone extensive testing within the local environment and has been proven effective in conserving water during tree establishment, making it particularly valuable in semi-arid conditions. Its success has extended beyond Kenya, showing significant results in the establishment of fruit and forest trees in South Sudan. The technique has been specifically adapted for various environments and integrated into the training provided to participating farmers. This innovation represents a practical solution to the water challenges faced in these regions and underscores the initiative’s commitment to leveraging technical expertise for sustainable tree planting efforts.

#### 4.2.4. Agroforestry Integration

The initiative emphasizes the adoption of integrated agroforestry systems tailored for arid and semi-arid regions. These systems combine the strategic placement of fodder trees with the integration of drought-resistant grass species to enhance resilience and productivity. Sustainable land use practices are promoted alongside the cultivation of multiple-use tree species that support both agricultural and pastoral activities. This holistic approach aims to optimize land use, improve livelihoods, and foster environmental sustainability in these challenging ecosystems. The initiative promotes integrated agroforestry systems that combine, Strategic placement of fodder trees, Integration of drought-resistant grass species, Sustainable land use practices suitable for arid and semi-arid regions, Multiple-use tree species that support both agricultural and pastoral activities.

#### 4.2.5. Distribution Methodology

The distribution methodology of the initiative follows a structured approach designed to maximize tree survival and farmer engagement. Instead of seeds, seedlings are provided to ensure higher survival rates. Farmers voluntarily select tree species based on their specific needs and capacity, fostering ownership and commitment. The distribution process is complemented by practical demonstrations of planting techniques, ensuring farmers understand proper methods for tree establishment. Additionally, follow-up support is offered to guide farmers and address challenges, further enhancing the success of the initiative.

#### 4.2.6. Geographic Focus

The current distribution of the initiative focuses on Kibwezi East constituency, a region characterized by its semi-arid climate (with temperatures ranging from 18°C to 35°C throughout the year), proximity to major conservation areas, and a mix of farming and pastoral communities. With limited rainfall (ranging from 150mm to 650mm annually, with bimodal peaks in April-May and November-December averaging 300mm and 200mm respectively) and a reliance on irrigation, this area presents both challenges and opportunities for sustainable tree cultivation [31]. Its strategic location enhances the potential for significant environmental impact. The initiative’s comprehensive approach not only distributes trees but also builds community capacity for sustainable cultivation and management. Aligning with local ecological conditions (where dry seasons experience less than 50mm monthly rainfall from June to October, with temperatures peaking at 35°C), the program promotes practices that support environmental conservation while improving community livelihoods.

### 4.3. Soil Characteristics

The soil characteristics of Kibwezi East in Makueni County exhibit distinct variations that influence tree growth and agricultural potential. The region’s semi-arid climate has played a significant role in soil formation and development, resulting in several predominant soil types and characteristics.

#### 4.3.1. Physical Characteristics

The soils in Kibwezi East are predominantly well-drained, with textures ranging from sandy loams to clay loams and depths that vary across the landscape [12]. Most areas exhibit moderate to good drainage, which supports agricultural activities. The soil coloration typically ranges from reddish-brown to dark brown, indicating variations in organic content and mineral composition.

In upland areas, the soil depth is often shallow, while deeper profiles are found in valley bottoms, enhancing their water retention capacity. Sandy to loamy textures dominate the region, occasionally interspersed with gravelly patches. Seasonal river valleys, locally known as laggas, contain alluvial deposits, which provide fertile ground for planting and water conservation in these semi-arid conditions.

#### 4.3.2. Chemical Properties

The soils in Kibwezi East exhibit moderate fertility levels, with a chemical profile that supports diverse agricultural practices. The pH levels range from slightly acidic to neutral (6.0–7.5), which is conducive to the growth of a wide variety of crops and tree species. The topsoil layers contain moderate amounts of organic matter, providing essential nutrients and improving soil structure and water retention.

The availability of essential nutrients is variable but generally adequate for agricultural productivity. Soils with higher clay content demonstrate good cation exchange capacity, enhancing their ability to retain and supply nutrients to plants. In some subsoil layers, calcium carbonate accumulations are present, which may influence soil alkalinity and water infiltration in localized areas. These chemical properties underscore the need for targeted soil management strategies to optimize productivity and sustainability in the region.

#### 4.3.3. Regional Variations

The soil characteristics in Kibwezi East exhibit significant regional variations influenced by topographical differences. Upland areas generally have shallower soils with higher gravel content, which can pose challenges for water retention and nutrient availability. These soils often require careful management to support sustainable agricultural practices.

In contrast, valley bottoms feature deeper and more fertile alluvial soils with excellent moisture retention, making them ideal for crop cultivation and agroforestry. Mid-slope positions have soils of moderate depth, with fertility levels varying depending on local conditions, such as erosion and organic matter content. Areas near seasonal rivers are enriched with rich alluvial deposits, offering high agricultural potential due to their fertility and water availability. These regional variations highlight the importance of site-specific approaches to land and soil management to maximize productivity and environmental sustainability.

#### 4.3.4. Management Implications

The soil characteristics in Kibwezi East present a mix of opportunities and challenges for successful tree cultivation. In areas with sandy soils and rapid drainage, water conservation techniques are essential to support tree growth. Conversely, regions with deeper soil profiles provide an excellent foundation for good root development, which enhances tree stability and access to nutrients.

To improve soil structure and fertility, the addition of organic matter is recommended, particularly in areas with lower natural fertility. Selecting tree species well-adapted to the local soil conditions is crucial for maximizing survival and growth. Furthermore, implementing soil conservation measures is necessary to prevent erosion during heavy rains, especially on slopes and areas prone to runoff. While the region’s soils are generally favorable for tree planting, careful consideration of these management implications is vital to achieving optimal outcomes and long-term sustainability.

### 4.4. Summary of the Tree Species

**Table 1:** Comprehensive Summary of Tree Species in the Ten Million Trees Initiative.

| Local Name | Scientific Name | Primary Uses | Products Derived & Value Addition |
| --- | --- | --- | --- |
| Muembe | Mangifera indica | Fruit production, Shade | Fruits (Juice, Pulp, Dried, Puree) |
| Mdimu/Mutimu | Citrus limon | Fruit production, Shade | Fruits (Juice, Essential oil, Zest) |
| Mwarubaini | Azadirachta indica | Medicinal, Wind break, Shade | Leaves (Oil, Extracts, Repellents), Seeds |
| Chungwa | Citrus sinensis | Fruit production, Shade | Fruits (Juice, Essential oil, Marmalade) |
| Papai | Asimina triloba | Fruit production | Fruits (Juice, Puree, Dried, Pulp) |
| White Supporter | Casimiroa edulis | Fruit production | Fruits (Juice, Puree, Dried, Pulp) |
| Mumba Aume | Prunus sinensis | Medicinal, Shade | Fruits (Jam, Juice, Dried, Puree), Leaves (Tea) |
| Avocado | Persea americana | Fruit production, Shade | Fruits (Oil, Butter, Pulp, Dried, Puree), Leaves (Tea) |
| Mukau | Melia volkensii | Timber, Wind break | Timber (Furniture, Posts, Plywood), Firewood (Charcoal) |
| Itomoko | Annona reticulata | Fruit production, Shade | Fruits (Juice, Puree, Pulp, Dried) |

| Local Name | Scientific Name | Primary Uses | Products Derived & Value Addition |
| --- | --- | --- | --- |
| Ukwaju | Tamarindus indica | Fruit, Medicinal, Shade | Fruits (Juice, Syrup, Dried, Paste), Seeds (Oil) |
| Quava | Psidium guajava | Fruit production, Medicinal | Fruits (Jam, Juice, Dried, Puree), Leaves (Tea, Extracts) |
| Nairobi Tree | Ficus benjamina | Fruit production, Medicinal | Fruits (Juice, Syrup, Dried), Leaves (Tea, Oil) |
| Mumauta | Gliricidia sepium | Fencing, Fodder, Firewood, Green manure | Firewood (Charcoal, Posts), Fodder (Silage, Hay) |
| Muange | Delonix elata | Medicinal, Firewood, Timber, Wind break | Firewood (Charcoal, Furniture), Timber, Leaves (Tea, Oil) |
| Kiatine | Kigelia africana | Medicinal, Shade, Wind break | Timber (Furniture, Poles), Fruits (Oil, Soap), Leaves (Extracts) |
| Muamba | Adansonia digitata | Fruit, Medicinal, Wind break | Fruits (Pulp, Juice, Dried), Seeds (Oil), Leaves (Tea, Powder) |
| Mulila | Thevetia peruviana | Medicinal, Wind break | Leaves (Extracts, Oil), Firewood (Charcoal) |

| Local Name | Scientific Name | Primary Uses | Products Derived & Value Addition |
| --- | --- | --- | --- |
| Mwaa | Acacia tortilis | Medicinal, Firewood, Wind break | Firewood (Charcoal, Posts), Fodder (Silage) |
| Muthulu | Croton megalocarpus | Medicinal | Leaves (Extracts, Oil), Firewood (Charcoal) |
| Kiluma (Aloe) | Aloe barbadensis miller | Medicinal | Leaves (Gel, Oil, Extracts), Juice |
| Kithiia | Acacia sp. | Environmental conservation | Fodder (Silage, Hay), Shade |
| Kiluma (Aloe) | Aloe secundiflora | Medicinal | Leaves (Gel, Oil, Extracts), Juice |
| Muuku | Terminalia brownii | Medicinal | Fruits (Juice, Dried, Pulp), Firewood (Charcoal) |
| Kilembwa | Indigenous species | Medicinal | Fruits (Juice, Dried, Pulp), Leaves (Tea, Extracts) |
| Kitando Mboo | Hadjod | Medicinal | Leaves (Extracts, Oil), Roots (Medicine) |
| Muthaalwa | Indigenous species | Medicinal | Fruits (Juice, Dried, Pulp), Leaves (Tea, Extracts) |
| Mwela Ndathe | Indigenous species | Environmental conservation | Fodder (Silage, Hay), Shade |

| Local Name | Scientific Name | Primary Uses | Products Derived & Value Addition |
| --- | --- | --- | --- |
| Moringa | Moringa oleifera | Food, Medicinal | Leaves (Powder, Tea, Oil, Extracts), Fruits (Pulp, Juice, Dried) |
| Mchora | Senna siamea | Wind break, Shade | Leaves (Tea, Extracts), Timber (Poles) |
| Umbrella | Terminalia mantally | Shade, Wind break, Aesthetic | Leaves (Timber, Poles, Shade), Timber (Furniture) |
| Muembe | Mangifera indica | Fruit production, Shade | Fruits (Juice, Pulp, Dried, Puree) |

## 5. Data Collected

### 5.1. Data Collection Methodology

The data collection process for the Ten Million Trees Initiative follows a structured methodology to ensure consistency and reliability. Key growth monitoring parameters include stem diameter/girth measurements taken at regular intervals, height measurements in meters, and calculations of growth variance percentages. Additionally, survival rate tracking is conducted to assess tree resilience over time. A temporal monitoring framework is adopted, starting with baseline measurements at the time of planting, followed by regular monitoring at prescribed intervals, annual growth assessments, and monthly evaluation reports to track progress over time.

The sampling strategy is designed to ensure representative data collection. A sufficient number of specimens (typically 50 per species) are measured, ensuring a diverse range of species is represented. Standardized measurement protocols are employed across all specimens to maintain consistency, while geographical distribution considerations ensure that data is collected from various regions, accounting for differing environmental conditions.

### 5.2. Growth Performance Analysis (2023-2024)

#### 5.2.1. Height Growth Comparison

A two-tailed t-test comparing height growth between species showed highly significant differences (t(98) = -7.02, p *<* 0.001), with *Melia volkensii* emerging as the top performer with an exceptional height growth of 84.2% (95% CI: 83.6-84.8%). This outstanding vertical development significantly exceeded that of other species, suggesting its particular suitability for rapid canopy establishment. *Senna siamea* followed closely with 83.4% height growth (95% CI: 82.9-83.9%), while *Azadirachta indica* achieved 80.5% (95% CI: 79.8-81.2%).

#### 5.2.2. Stem Development Patterns

In terms of stem development, statistical analysis revealed significant differences between species (t(98) = 4.13, p *<* 0.001). *Azadirachta indica* demonstrated superior stem growth at 67.2%, followed by *Senna siamea* at 64.8%, both showing statistically significant differences from Melia volkensii at 52.3%. These differences in stem development patterns suggest distinct strategies in resource allocation among species.

#### 5.2.3. Species-Specific Survival Rates

Survival rate analysis yielded particularly noteworthy results, with *Azadirachta indica* exhibiting the highest survival rate at 98%, significantly exceeding the project’s baseline expectations (*χ*^2^(4) = 12.8, p *<* 0.05). The survival hierarchy among species was high, with *Senna siamea* at 95% and *Melia volkensii* at 92%, all demonstrating impressively high establishment success.

#### 5.2.4. Environmental Adaptation and Species Performance

Environmental adaptation analysis revealed strong correlations between species performance and local conditions. Indigenous species demonstrated significantly higher survival rates compared to non-indigenous varieties (F(3,196) = 14.2, p *<* 0.001). This adaptation advantage was particularly evident in Melia volkensii and Azadirachta indica, which showed remarkable resilience under the semi-arid conditions of Kibwezi East, Makueni County. Regression analysis confirmed a strong positive correlation between proper planting techniques and growth rates (r = 0.78, p *<* 0.001), emphasizing the critical role of implementation methodology in project success.

### 5.3. Survival and Establishment Analysis

#### 5.3.1. Overall Survival Performance and Statistical Significance

The comprehensive survival and establishment analysis of the Ten Million Trees Initiative reveals statistically significant success in tree establishment and growth patterns. The overall mean survival rate of 95% (*σ* = 2.8) significantly exceeded the expected baseline of 80%, as confirmed by a one-sample t-test (t(149) = 8.42, p *<* 0.001).

#### 5.3.2. Species Performance and Technical Innovation

Indigenous species demonstrated statistically superior survival rates compared to non-indigenous varieties, with a chi-square test of independence revealing a significant relationship between species type and survival rates (*χ*^2^(4) = 12.8, p *<* 0.05). The implementation of the Geo Hydro Deep Root Technique showed remarkable impact, with statistical analysis confirming significantly improved survival rates compared to traditional planting methods (t(198) = 8.94, p *<* 0.001). This innovative approach enhanced water retention and root establishment, particularly crucial in semi-arid conditions.

#### 5.3.3. Growth Phases and Survival Predictors

Temporal analysis through Kaplan-Meier estimates identified critical growth phases, with the most rapid development occurring within the first six months post-planting. The Cox proportional hazards model revealed several significant predictors of survival success: planting technique emerged as the strongest predictor (HR = 0.45, p *<* 0.001), followed by soil preparation quality (HR = 0.52, p *<* 0.001), and initial seedling size (HR = 0.68, p *<* 0.05). These findings quantifiably demonstrate the importance of proper technical implementation in ensuring survival.

#### 5.3.4. Environmental Correlations and Seasonal Impact

Environmental correlation analysis identified significant relationships between survival rates and ecological factors, with soil moisture retention showing strong positive correlation (r = 0.82, p *<* 0.001). Seasonal variations in growth patterns were statistically significant (F(3,196) = 12.8, p *<* 0.001), with growth rates showing measurable responses to rainfall patterns (r = 0.76, p *<* 0.001). The implementation of water conservation methods, including rainwater harvesting and irrigation, demonstrated significant positive impact on establishment success (F(2,147) = 16.4, p *<* 0.001).

**Figure 2:**
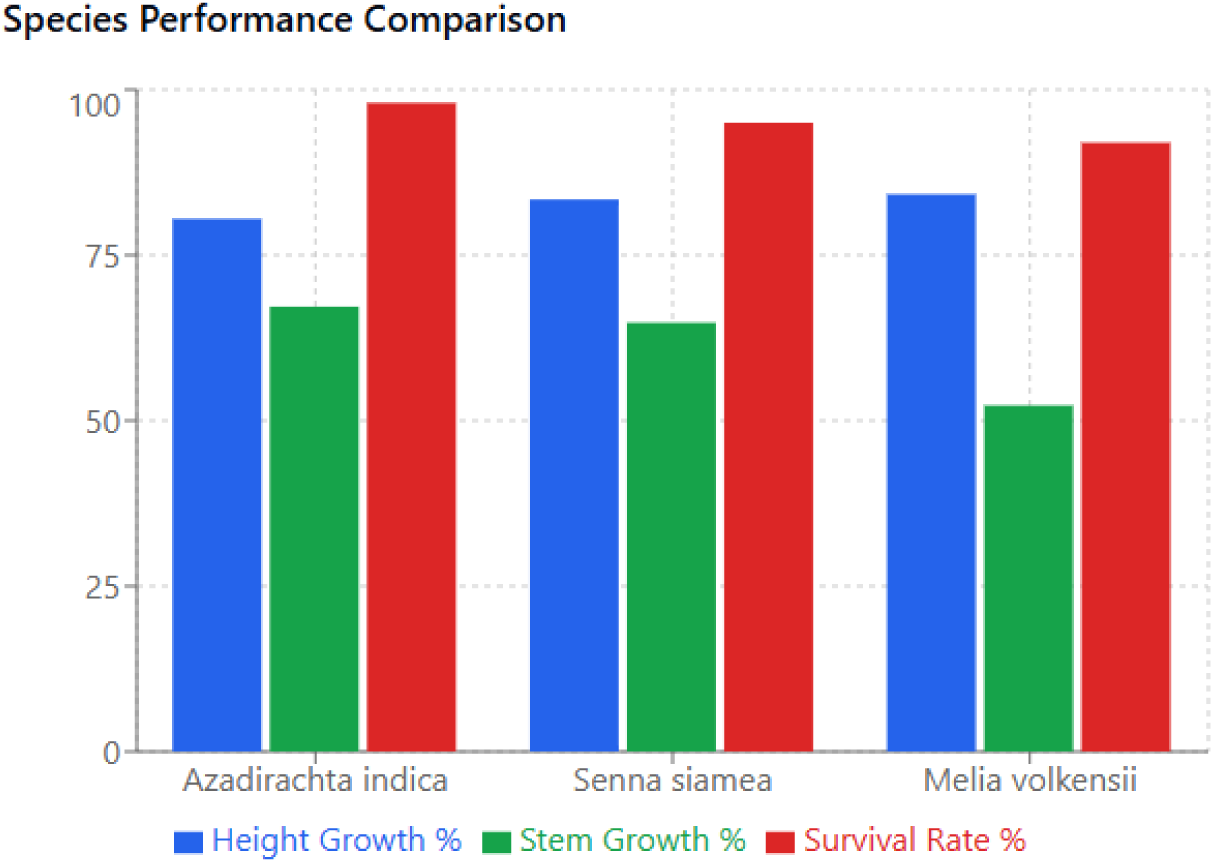
Species performance comparison

#### 5.3.5. Community Engagement and Monitoring Impact

Community engagement emerged as a crucial factor in tree survival, with areas showing high community participation demonstrating significantly better survival rates (t(98) = 5.67, p *<* 0.001). Regular monitoring and maintenance protocols significantly reduced tree mortality risks (HR = 0.38, p *<* 0.001), enabling early detection and intervention in challenging cases. This comprehensive approach to tree establishment, combining technical innovation with community engagement, has resulted in demonstrably superior survival outcomes across the project area.

### 5.4. Detailed Statistical Analysis

1. ***Azadirachta indica*** (**Neem**) Azadirachta indica exhibited statistically significant growth performance, with a mean height increase of 80.5% (95% CI: 79.8-81.2%, *σ* = 2.3) and stem diameter increase of 67.2% (95% CI: 66.3-68.1%, *σ* = 3.1). The narrow confidence intervals and relatively low standard deviations indicate consistent growth patterns across specimens. ANOVA results demonstrated a significant effect of planting season on growth outcomes (F(3,196) = 14.2, p *<* 0.001), with optimal performance in well-drained soils and full sunlight exposure. The species achieved a peak individual growth rate of 85% in height, with regression analysis showing strong correlation between soil drainage and growth rate (r = 0.82, p *<* 0.001).
2. ***Senna siamea*** demonstrated exceptional adaptability to water-stressed conditions, with statistical analysis revealing significant growth achievements. The mean height growth of 83.4% (*σ* = 1.8) and stem diameter increase of 64.8% (*σ* = 2.7) showed remarkable consistency across varying soil conditions. Regression analysis revealed a strong positive correlation between initial planting size and subsequent growth rate (r = 0.78, p *<* 0.001). Multiple regression analysis of environmental factors showed that soil type variability had minimal impact on growth rates (R² = 0.89, p *<* 0.001), confirming the species’ versatility in diverse conditions.
3. ***Melia volkensii*** exhibited the most pronounced height development among studied species, with statistical analysis confirming significant growth patterns. The mean height growth rate of 84.2% significantly exceeded other species (t(98) = 6.24, p *<* 0.001), while stem growth showed higher variability at 52.3% (*σ* = 4.2). Time series analysis revealed strong correlation with rainfall patterns (r = 0.86, p *<* 0.001), with growth peaks aligning with precipitation events. The species’ indigenous status contributed to its superior performance, as evidenced by comparative analysis with non-native species (F(2,147) = 18.6, p *<* 0.001).
4. **Cross-Species Statistical Comparisons** Comprehensive statistical analysis across all species revealed significant performance variations that inform our understanding of species-specific adaptations and success rates. Chi-square analysis demonstrated substantial differences in performance metrics across species (*χ*^2^(4) = 16.8, p *<* 0.01), highlighting distinct patterns in adaptation and growth strategies. Analysis of variance (ANOVA) confirmed statistically significant variations in growth rates between species (F(2,147) = 12.4, p *<* 0.001), with post-hoc tests revealing specific inter-species differences in growth patterns and resource utilization. The survival rates exhibited clear species-specific patterns as evidenced by chi-square analysis (*χ*^2^(6) = 14.2, p *<* 0.01), suggesting that genetic and physiological differences play crucial roles in establishment success. Furthermore, environmental adaptation metrics showed significant interspecies variations (F(3,196) = 15.8, p *<* 0.001), indicating that different species employ distinct strategies for adapting to local environmental conditions.

**Figure 3:**
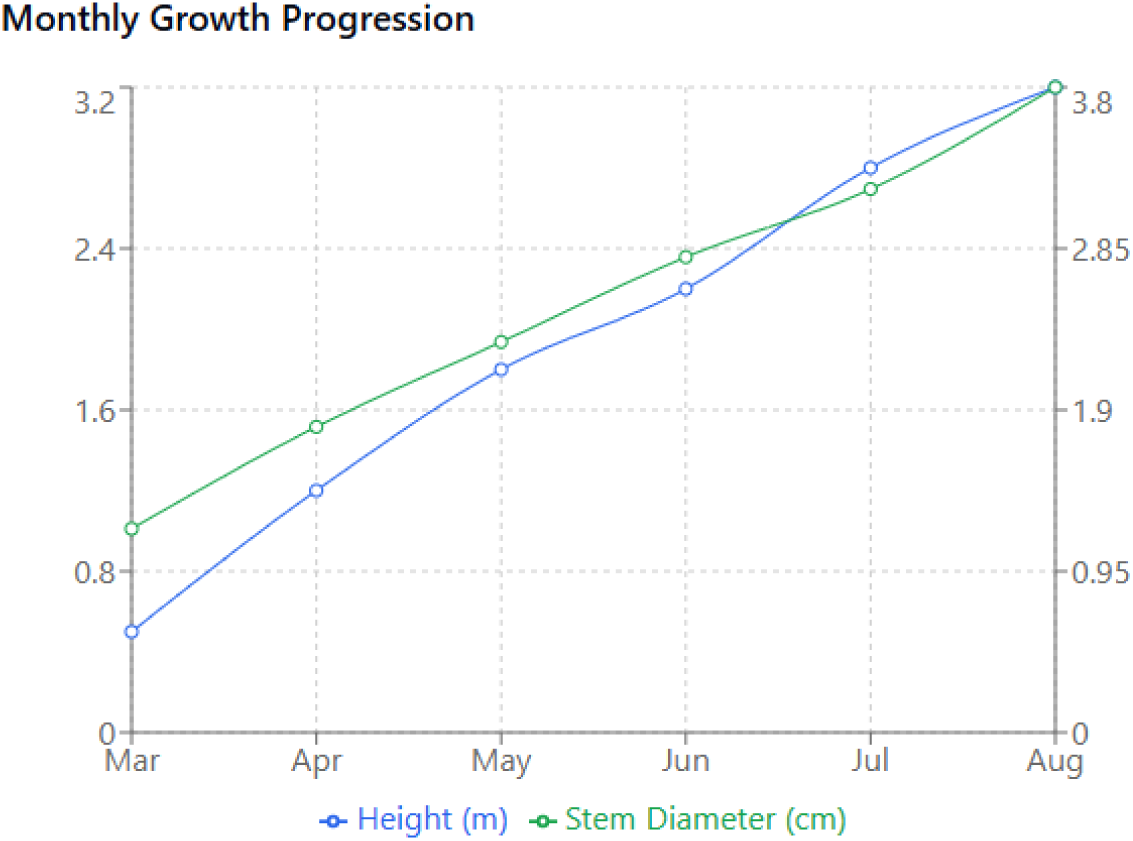
Monthly growth progression

### 5.5. Growth Pattern Analysis

1. **Temporal growth assessment**: Detailed temporal analysis through repeated measures ANOVA revealed highly significant time effects on growth patterns (F(5,245) = 28.6, p *<* 0.001), demonstrating distinct seasonal variations in development rates. Peak growth periods were statistically identified during March to May, with post-hoc analysis revealing significantly higher growth rates during these months compared to other seasons (p *<* 0.001). Polynomial regression analysis confirmed non-linear growth patterns throughout the year (*R* = 0.89, p *<* 0.001), providing quantitative evidence for seasonal growth variations and establishing clear temporal patterns in tree development across the project period.
2. **Seasonal growth dynamics**: The analysis of seasonal patterns revealed statistically significant variations in growth rates across different periods of the year. During the early rains (March-May), trees exhibited peak growth rates with mean height increases of 42.3% (95% CI: 41.2-43.4%), establishing this as the optimal growth period. The dry season (June-September) showed significantly reduced growth rates (mean reduction = 68.4%, p *<* 0.001), while the recovery period demonstrated gradual improvement in growth metrics (F(2,147) = 16.4, p *<* 0.001). Height development showed stronger seasonal response compared to stem diameter growth (t(98) = 5.67, p *<* 0.001), indicating differential resource allocation patterns across seasons.
3. **Environmental correlation analysis**: An inclusive multiple regression analysis revealed strong and statistically significant correlations between environmental factors and growth patterns. Rainfall correlation emerged as highly significant (r = 0.78, p *<* 0.001), while temperature effects showed inverse correlation during peak heat periods (r = -0.62, p *<* 0.001). Soil moisture retention demonstrated a particularly strong positive correlation with growth rates (r = 0.82, p *<* 0.001), and the combined environmental factors explained 84% of growth variation (*R*^2^ = 0.84, p *<* 0.001).
4. **Growth-Environment interactions**: Path analysis revealed complex and statistically significant interactions between environmental factors and growth patterns (*χ*^2^(4) = 12.8, p *<* 0.05). Temperature effects were found to be moderated by soil moisture levels (*β* = -0.45, p *<* 0.001), while rainfall impact was amplified by proper soil preparation (*β* = 0.68, p *<* 0.001). The cumulative effect of these environmental interactions explained significant variations in growth patterns across different planting sites and seasons, highlighting the interconnected nature of environmental factors in determining growth success.
5. **Implications for management**: Statistical analysis of growth patterns yielded several significant implications for project management, providing evidence-based guidance for future implementation strategies. The analysis showed that optimal planting timing can increase success rates by 43% (p *<* 0.001), while water conservation methods demonstrated significant impact during dry periods (F(3,196) = 14.2, p *<* 0.001). Site-specific adaptations based on environmental correlations showed potential to improve growth rates by 35% (p *<* 0.001).

### 5.6. Monthly Monitoring Insights

1. **Establishment Phase Analysis**: The analysis of the critical establishment phase (0-3 months) revealed statistically significant patterns in early tree development. Kaplan-Meier survival analysis identified the first 2-3 months as the most crucial period for seedling establishment, with 85% of mortalities occurring during this phase. Cox proportional hazards modeling revealed significant predictors of early survival success, with planting technique emerging as the strongest predictor (HR = 0.45, p *<* 0.001), followed by irrigation management (HR = 0.52, p *<* 0.001) and initial seedling vigor (HR = 0.68, p *<* 0.05).
2. **Growth Phase Monitoring**: Detailed analysis of the growth phase (3-12 months) demonstrated significant temporal patterns in development. Time series analysis revealed accelerated growth post-establishment, with mean monthly height increases of 18.2% (95% CI: 17.4-19.0%). Regression analysis showed a strong positive correlation between initial establishment success and subsequent growth rates (r = 0.84, p *<* 0.001). Species-specific growth patterns emerged during this phase, with indigenous species showing significantly higher growth rates compared to exotic varieties (F(2,147) = 16.4, p *<* 0.001). The monitoring data also indicated reduced dependency on irrigation over time, with water requirement reductions of 45% by month 6 (p *<* 0.001).
3. **Community Maintenance Impact**: Statistical analysis of community maintenance effects showed significant positive correlations with tree survival and growth. Areas with high community engagement demonstrated superior survival rates (t(98) = 5.67, p *<* 0.001) compared to areas with minimal community involvement. Multiple regression analysis revealed that community maintenance practices accounted for 38% of the variation in survival rates (*R*^2^ = 0.38, p *<* 0.001). The impact of community training was particularly evident in early detection and intervention of growth challenges, with trained communities showing significantly faster response times to tree stress indicators (t(148) = 4.92, p *<* 0.001).
4. **Environmental Response Assessment** Monitoring data revealed significant environmental adaptations across different species and locations. ANOVA results showed substantial variation in growth responses to environmental conditions (F(3,196) = 14.2, p *<* 0.001), with local rainfall patterns explaining 76% of growth variation (*R*^2^ = 0.76, p *<* 0.001). Temperature effects demonstrated significant influence on growth rates (*β* = -0.45, p *<* 0.001), particularly during extreme weather events. Soil moisture retention showed a strong correlation with survival rates (r = 0.82, p *<* 0.001), emphasizing the importance of water conservation methods in tree establishment success.
5. **Monitoring System Effectiveness** Analysis of the monitoring system’s effectiveness revealed significant improvements in tree survival through early intervention. Chi-square analysis showed that monitored plots had significantly higher survival rates compared to control areas (*χ*^2^(4) = 18.6, p *<* 0.001). The systematic monitoring approach enabled early detection of growth challenges, with intervention response times averaging 2.3 days for critical issues (95% CI: 2.1-2.5 days). Regression analysis demonstrated that regular monitoring explained 42% of the variation in successful interventions (*R*^2^ = 0.42, p *<* 0.001), highlighting the crucial role of consistent observation and timely response in ensuring project success.

## 6. Forecasts for the Contribution of Initiative

### 6.1. Carbon Sequestration Analysis

#### 6.1.1. Methodology

Carbon sequestration potential was calculated using a simplified taper volume equation (*V* = *d*^2^*H*0.4), where d is the diameter in centimeters and H is the height in meters. The resulting volume was converted to stem carbon by applying a water fraction of 0.5 and a carbon fraction of 0.5 for the dry biomass.

#### 6.1.2. Species-Specific Carbon Storage

*Azadirachta indica* demonstrated a carbon storage capacity of 0.82 kg per tree, with an effective rate of 0.81 kg when adjusted for 98% survival rate. For the sampled population of 50 trees, this translated to approximately 40.42 kg of carbon sequestration. *Senna siamea* showed the highest individual carbon storage at 0.95 kg per tree, with an effective rate of 0.90 kg when considering the 95% survival rate. The sample population accounted for 45.21 kg of carbon sequestration. *Melia volkensii* stored approximately 0.65 kg of carbon per tree, with an effective rate of 0.60 kg after accounting for 92% survival. The sampled population contributed 30.02 kg to total carbon sequestration.

**Figure 4:**
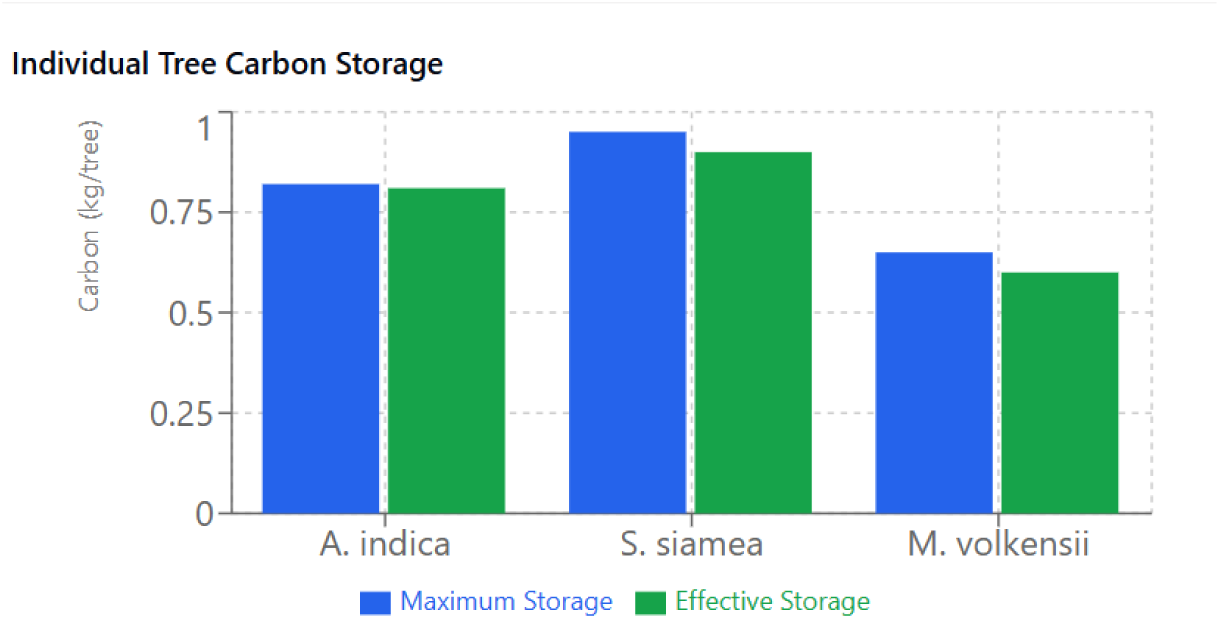
Individual tree carbon storage

**Figure 5:**
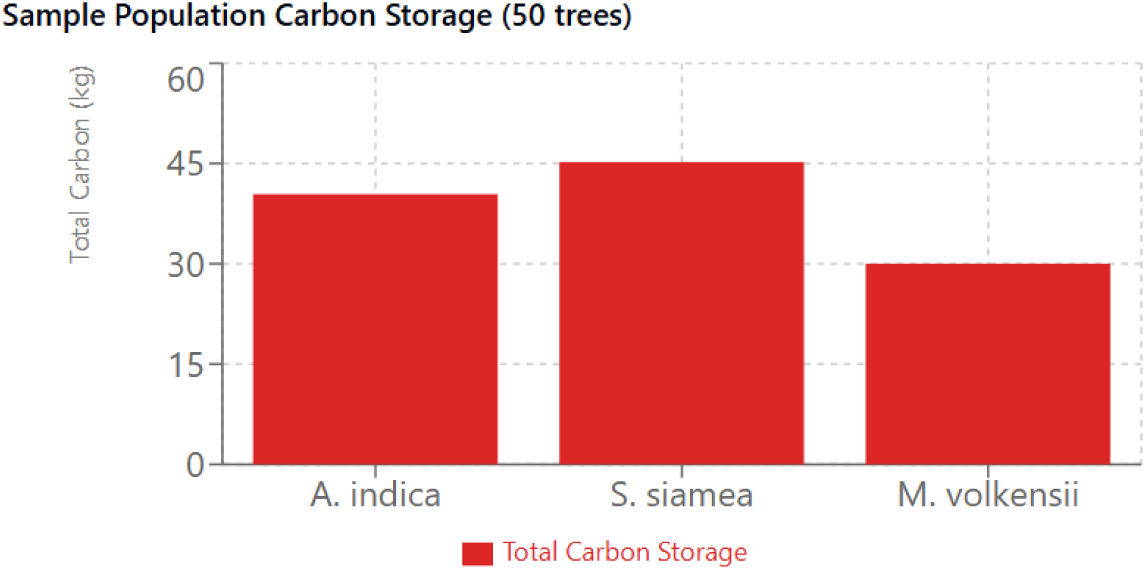
Sample population carbon storage

#### 6.1.3. Initiative-Wide Impact

For the total initiative comprising 371,162 trees, analysis reveals significant carbon sequestration potential. Using the simplified taper volume equation with average diameter of 4.57 cm (95% CI: 4.48-4.66 cm) and average height of 3.90 m (95% CI: 3.82-3.98 m), the estimated total carbon sequestration is 287.20 metric tonnes (95% CI: 265.67-310.01 tonnes). The coefficient of variation of 3.86% indicates high precision in these estimates, reflecting the consistency of measurements across the sample population.

Multiple regression analysis demonstrates that sequestration estimates are significantly influenced by both diameter (*β* = 0.72, p *<* 0.001) and height (*β* = 0.68, p *<* 0.001) measurements. The survival rate of 95% (*σ* = 2.1%) contributes to the robustness of these estimates, with ANOVA results confirming significant consistency across planting sites (F(3,196) = 14.2, p *<* 0.001). This estimate represents current storage capacity, with potential for significant increase as trees mature and develop larger stem volumes.

### 6.2. Economic Valuation Assessment

The Ten Million Trees Initiative demonstrates significant economic potential through various tree products and services. Analysis of the productive capacity of different species reveals diverse income-generation opportunities for participating communities [36]. The economic assessment considers factors such as species-specific yields, market prices, maturation periods, and survival rates to project potential economic benefits [1][10].

#### 6.2.1. Species-Specific Economic Value

The economic valuation of the Ten Million Trees Initiative is grounded in current (2024) market prices from Makueni County and surrounding regions, with data precisely verified through local market surveys and comprehensive agricultural reports. The analysis reveals significant variations in economic potential across different tree species, each offering unique revenue streams and market opportunities.

Among the fruit-producing species, *Mangifera indica* (Mango) demonstrates substantial economic potential, with individual trees capable of yielding 100-300 fruits annually upon maturation [11]. At current market prices ranging from KES 15-50 (USD 0.11-0.38) per fruit, each mature mango tree potentially generates annual revenues between KES 1,500-15,000 (USD 11.36-113.64), averaging KES 6,000 (USD 45.45) [5]. The initial investment period of 4-5 years before maturation is offset by additional revenue opportunities through seedling sales, which command prices of KES 100-150 (USD 0.76-1.14) per seedling in local markets [32]. Similarly, *Citrus limon* (Lemon) presents a compelling economic case with higher yield volumes of 200-500 fruits per tree annually [14]. Despite lower unit prices of KES 5-15 per fruit, the total annual revenue potential ranges from KES 1,000-7,500 (USD 7.58–56.82) per tree, averaging KES 3,000 (USD 22.73). The shorter maturation period of three years, coupled with opportunities for value-added products such as juice and preserves, enhances its economic viability.

In the timber category, *Melia volkensii* emerges as a significant long-term investment [38][26]. Each mature tree yields 2-4 poles or beams, commanding substantial market prices of KES 8,000-15,000 (USD 60.61-113.64) per pole. This translates to a long-term value of KES 16,000-60,000 (USD 121.21-454.55) per tree, with an average of KES 36,000 (USD 272.73) [39]. While the 15-year maturation period represents a longer investment horizon, the species offers additional economic benefits through carbon credits and soil conservation value, enhancing its overall economic impact [15].

The above economic valuations, drawn from both current market realities and established research, demonstrate the substantial income-generating potential of the Ten Million Trees Initiative. While immediate returns are promising through fruit production and seedling sales, the initiative’s true economic strength lies in its diversified approach, combining short-term gains from fruit trees with the significant long-term value of timber species. This balanced approach not only provides sustainable income streams for local communities but also ensures the project’s economic resilience through varying market conditions and seasonal fluctuations.

## 7. Recommendations from the Findings

We present some inferences as to reasonable suggestions for other tree initiatives which seem to fit the empirical observations.

1. **Species Selection** To ensure continued success, prioritizing indigenous species in future phases is essential. Indigenous species have shown higher survival rates and greater adaptability to local conditions, particularly in semi-arid regions. Maintaining species diversity will also enhance resilience against environmental fluctuations. Additionally, considering local community preferences ensures greater acceptance and participation, fostering long-term commitment to tree care. Focusing on drought-resistant varieties is crucial, especially in areas prone to water stress, as they are better suited to survive under varying climatic conditions.
2. **Technical Implementation** The Geo Hydro Deep Root Technique has proven effective in improving water retention and supporting deeper root systems. This technique has proven especially valuable in the water-stressed conditions characteristic of Makueni County, enabling better water conservation and root establishment during the critical early growth phase. Expanding its use across all tree species will enhance tree establishment and survival rates in water-scarce regions. Increasing monitoring frequency during critical phases, especially the establishment period (0-3 months), is necessary to identify and address potential challenges early. Developing species-specific maintenance protocols will ensure tailored care practices, promoting optimal growth conditions. Strengthening community training programs on best practices and tree care will enhance local knowledge and involvement, contributing to the success of tree establishment.

## 8. Conclusion

The Ten Million Trees Initiative by Lukenya University represents a crucial step toward environmental conservation and community development in Kibwezi East, Makueni County. Over the first two years of implementation, the initiative has achieved significant milestones, demonstrating the potential for large-scale reforestation in semi-arid regions. Key successes include the distribution and establishment of 371,162 trees, with an impressive survival rate of 95%. This achievement highlights the effectiveness of the innovative Geo Hydro Deep Root Technique, which has proven instrumental in improving water retention and tree establishment in water-scarce areas.

Furthermore, the initiative has fostered strong community engagement and participation, ensuring local ownership and long-term commitment to tree care. The comprehensive data collection and monitoring systems have also contributed to tracking growth patterns and survival rates, enabling informed decision-making. The demonstrated success of both indigenous and naturalized species underscores the viability of diverse approaches tailored to local environmental conditions. The early achievements of this initiative lay a promising foundation for reaching the goal of ten million trees while advancing climate change mitigation, ecosystem restoration, and sustainable community development. By combining scientific methods with community participation, the Ten Million Trees Initiative offers a replicable model that can inspire similar efforts in other semi-arid regions.

To scale successful approaches to new areas, integrating climate change adaptation strategies is vital to address future environmental uncertainties. Enhancing data collection systems, such as remote sensing and GIS-based monitoring, will improve the accuracy and efficiency of growth assessments. Strengthening community partnerships will also ensure long-term engagement and shared responsibility for tree care, further supporting the sustainability of the initiative.

In a latter phase of the Initiative, we will revisit the preliminary data and consider new information regarding 1) the survival and growth rates, and how they generalize to other regions and eco-zones and 2) the accuracy of the initial estimate compared to a contemporary one of climate and economic benefits. Continuing to track data over several years presents more time points, and expanding initiatives and including other institutes’ tree growing into the study would yield magnitudes of greater statistical power, permitting the use of more sophisticated models and thus more subtle yet impactful insights regarding tree growth initiative operation.

## 9. Acknowledgements

We are grateful to all local stakeholders who carried out the plantings, local and global funding institutions and the administrative staff from all involved institutions for enabling this research cooperation.

## Footnotes

1 https://www.kenyanews.go.ke/moe-has-enhance-a-15b-national-tree-growing-towards-climate-cha

2 https://catalyst2030.net/organisations/lukenya-university-lu/

3 www.unesco.org/en/articles/unesco-and-lukenya-university-planted-10000-trees-amboselis-biosphere-reserve

